# The mRNA translation landscape in the synaptic neuropil

**DOI:** 10.1101/2020.06.09.141960

**Authors:** Caspar Glock, Anne Biever, Georgi Tushev, Ina Bartnik, Belquis Nassim-Assir, Susanne tom Dieck, Erin M. Schuman

## Abstract

To form and modify synaptic connections and store information, neurons continuously remodel their proteomes. The impressive length of dendrites and axons imposes unique logistical challenges to maintain synaptic proteins at locations remote from the transcription source (the nucleus). The discovery of thousands of mRNAs near synapses suggested that neurons overcome distance and gain autonomy by producing proteins locally^1^. It is not known, however if, how and when localized mRNAs are translated into protein. To investigate the translational landscape in neuronal subregions, we performed simultaneous RNA-seq and Ribo-seq from microdissected rodent brain slices to identify and quantify the transcriptome and translatome in cell bodies as well as dendrites and axons (neuropil). More than 4800 transcripts were translated in synaptic regions. Thousands of transcripts were differentially translated between somatic and synaptic regions, with scaffold and signaling molecules mostly arising from local sources. Furthermore, specific mRNA features were identified that regulate the efficiency of mRNA translation. The findings overturn the view that local translation is a minor source of synaptic protein^2^ and indicate that on-site translational control is an important mechanism to control synaptic strength.

## Main text

At neuronal synapses, more than 2500 proteins^1,3^ (the “synaptic proteome”) act as sensors and effectors to control neuronal excitability, synaptic strength and plasticity. The elaborate morphology and functional compartmentalization of the individual neuron imposes unique logistical challenges to maintain and modify the synaptic proteome at locations remote from the transcription source (i.e. the nucleus). To fulfill the local demand for new protein, neurons localize messenger RNAs (mRNAs) and ribosomes near synapses to produce proteins directly where they are needed^1^. Using high-throughput sequencing, several groups have reported the localization of thousands of transcripts to axons and dendrites (the “local transcriptome”)^4-8^. In many cell types, however, it has been shown that the transcript levels do not always predict protein levels^9^, suggesting that mRNA translation is a highly regulated process. Since proteins, rather than mRNAs, drive cellular function, we must therefore determine directly which transcripts are translated into proteins in dendrites and/or axons in vivo (the “local translatome”). Importantly, the relative contribution of a central (soma) versus local (dendrites/axons) source for individual proteins also remains unknown.

A given transcript’s translation level is determined by the rate of ribosome recruitment to the start codon during initiation and the velocity of ribosome translocation during polypeptide elongation. For most mRNAs, translation initiation is considered rate-limiting^10^: Initiation is regulated by elements within the mRNA’s untranslated regions (UTRs) that bind RNA-binding proteins (RBPs) or miRNAs^11-13^. In addition, the elongation rate also plays a regulatory role in determining the amount of protein produced from a transcript^14^. Although disrupted translational control has been linked to a number of neurological disorders^15^, little is known about the magnitude and mechanisms for transcript-specific translational regulation in neuronal compartments.

In this study, we combined deep-sequencing of ribosome-protected fragments (Ribo-seq) and RNA-seq on microdissected hippocampal rodent brain sections to provide a comprehensive analysis of the mRNA translational landscape both in cell bodies and the neuropil (a region enriched in neuronal dendrites/axons). We identified more than 4800 mRNAs that were translated in the synaptic neuropil and validated their local translation using high-resolution visualization of newly synthesized protein *in situ*. We found evidence for pervasive translational regulation of synaptic proteins in neuronal processes. These results reveal an unprecedented capacity for local protein production in vivo to maintain and modify the pre- and post-synaptic proteome.

### More than 4800 mRNAs are translated in synaptic regions

To discover the mRNA species localized and translated in cell-bodies as well as dendrites and axons we carried out a genome-wide analysis of the somata and neuropil transcriptome and translatome, using a dataset from microdissected hippocampal slices^16^. Ribosome footprints were obtained from somata and neuropil lysates to assess the number and position of translating ribosomes on a transcript (Ribo-seq^17^). In parallel, transcript levels were quantified by performing RNA-seq from the somata and neuropil (Fig. 1a). RNA- and Ribo-seq libraries from the somata and neuropil were highly reproducible among the three biological replicates (Fig. S1a, b). Furthermore, the Ribo-seq samples exhibited the expected depletion of footprint read densities in the UTRs and introns of transcripts (Fig. S1c, d), as well as three-nucleotide phasing (Fig. S1e, f)^17^.

**Figure 1.**
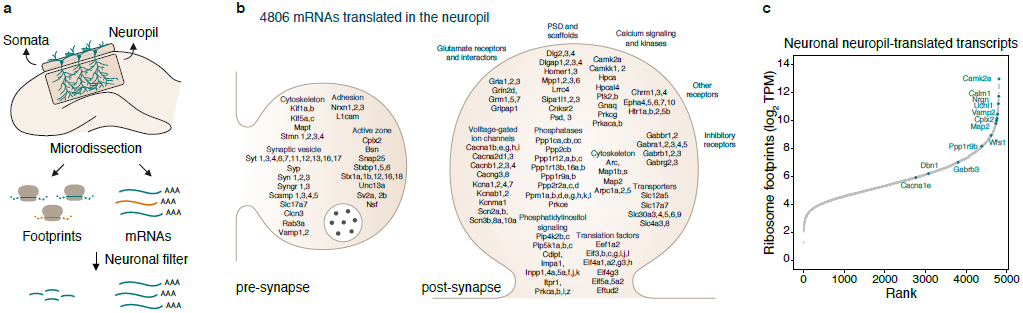
More than 4800 mRNAs undergo translation in dendrites and/or axons. (**a**) Experimental workflow. Microdissection of the CA1 region of the rat hippocampus. RNA-seq and Ribo-seq were conducted simultaneously for the somata (enriched in pyramidal neuron cell bodies) and the neuropil (enriched in dendrites and axons) layers. A neuronal filter was applied to enrich for excitatory neuron transcripts in downstream analyses. (**b**) Scheme of the pre- and post-synaptic compartments highlighting some of the 4806 locally translated transcripts highly relevant for synaptic function. (**c**) Rank plot showing locally translated transcripts sorted by ribosome density (log_2_ TPM). Selected synaptic transcripts are highlighted in teal.

The somata and neuropil of the hippocampus contain excitatory neuron cell bodies and their processes, as well as glia and interneurons. We created a pipeline to focus on excitatory neuron genes by minimizing the contribution of other cell types via bioinformatic filtering (Fig. S2a). To obtain a comprehensive set of glia-enriched transcripts, we prepared hippocampal neuron- and glia-enriched cultures (Fig. S2b; see Methods). In addition, we identified the transcripts de-enriched in excitatory neurons from the hippocampus (see Methods). Combining these datasets, we obtained a list of “contaminant” non-excitatory neuron genes (Fig. S2a, c) that was then subtracted from the somata or neuropil Ribo-seq datasets to generate and vet^18^ a list of 5521 excitatory neuron transcripts (Fig. S2a, d).

We identified the population of excitatory-neuron transcripts that undergo translation in the neuropil of the hippocampus. We detected with high-fidelity the translation of 4806 mRNAs, (the local translatome), in the neuropil (Fig.1b, Fig. S2a, see Methods). Within the local translatome we found many mRNAs encoding proteins that function in pre- and post-synaptic compartments, including scaffold molecules (e.g. *Dlg4, Homer1, Dlgap1*), receptors (e.g. *Gria1, Grin2d, Grm5*), ion channels (e.g. *Cacna1b, Kcnab2, Scn2a*), signaling molecules (e.g. *Camk2a, Hpca, Ppp1r9a*), synaptic vesicle (e.g. *Slc17a7, Syp*) and active zone proteins (e.g. *Bsn*) (Fig. 1b). Because each ribosome engaged with an mRNA produces one protein molecule, the translation level of a given transcript is proportional to its ribosome density (measured by the footprint number per unit length of the mRNA (see Methods)^17,19^. A ranking of ribosome densities of mRNAs in the neuropil revealed that many transcripts encoding key regulators of synaptic strength, such as *Camk2a* or *Nrgn*, were amongst the most highly translated in the neuropil (Fig. 1c).

To validate the local translation of the transcripts identified in the neuropil Ribo-seq data, we visualized newly synthesized target proteins in situ using Puro-PLA, a puromycin-based labeling strategy^20^. Puro-PLA enables the coincident detection of a protein of interest and a metabolic label in neurons (Fig. 2a). Cultured hippocampal neurons were briefly (5 min) incubated with puromycin^21^, then processed to identify nascent proteins-of-interest encoded by the local translatome mRNAs (see Methods). Indeed, for all of the “local translatome” proteins we assessed, we detected considerable nascent protein signal in dendrites (Fig. 2b), thus validating the Ribo-seq translatome data. We also confirmed the specificity of the nascent protein signal by its substantial reduction when puromycin was omitted (Fig. S3) or when neurons were pre-treated with the protein synthesis inhibitor anisomycin (Fig. S3). Taken together, our data reveal a surprisingly high number of protein species that are synthesized in the synaptic neuropil in vivo.

**Figure 2.**
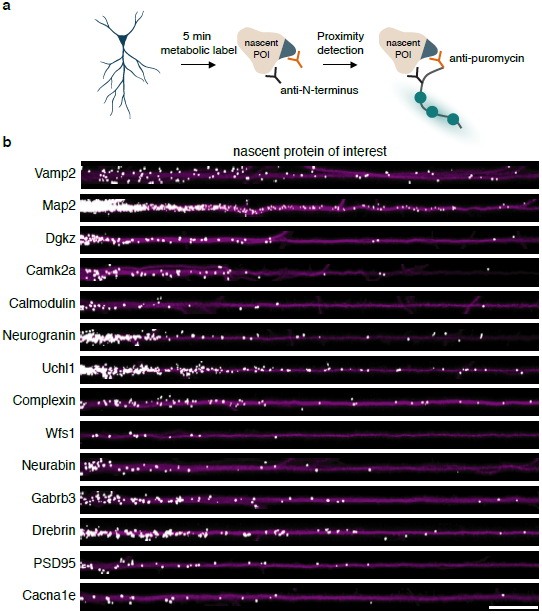
Validation of synthesis of proteins-of-interest in neuronal processes by high-resolution fluorescent imaging in situ. (**a**) Experimental workflow. Nascent proteins of interest (POI) were metabolically labeled with puromycin for 5 min and visualized in hippocampal neurons using proximity detection (Puro-PLA^20^). (**b**) Puro-PLA signal of candidate POI as indicated (white) in individual straightened dendrites from cultured hippocampal neurons. Signal was dilated for better visualization. Anti-MAP2 immunolabeling of dendrites is indicated in magenta. Scale bar = 20 *µ*m.

### Pervasive translational regulation of synaptic proteins in dendrites and axons

To investigate if transcripts in dendrites and/or axons are translationally regulated, we compared neuropil RNA and ribosome footprint levels for each transcript. As expected, the neuropil ribosome footprints and mRNA levels generally correlated well, with more abundant mRNAs displaying increased ribosome footprint abundance (R^2^ = 0.705; Fig. 3a). Some mRNAs, however, exhibited a weaker correlation between transcript and translation levels, indicating positive or negative translational regulation. To measure directly the translational efficiency (TE) of neuropil transcripts, we computed the ratio of ribosome footprints (from Ribo-seq) to mRNA fragments (from RNA-seq)^17^. We observed a wide distribution of translation efficiencies, with a greater than 200-fold difference between the most and least efficiently translated transcripts in the neuropil (Fig. 3b). In particular, we identified 892 and 896 transcripts exhibiting a significantly higher (TE_high_, e.g. *Camk2a* and *Eef1a2*, Fig. 3c) or lower (TE_low_, e.g. *Kif5c* and *Homer2*, Fig. 3c) TE, respectively (Fig. 3b, see Methods).

**Figure 3.**
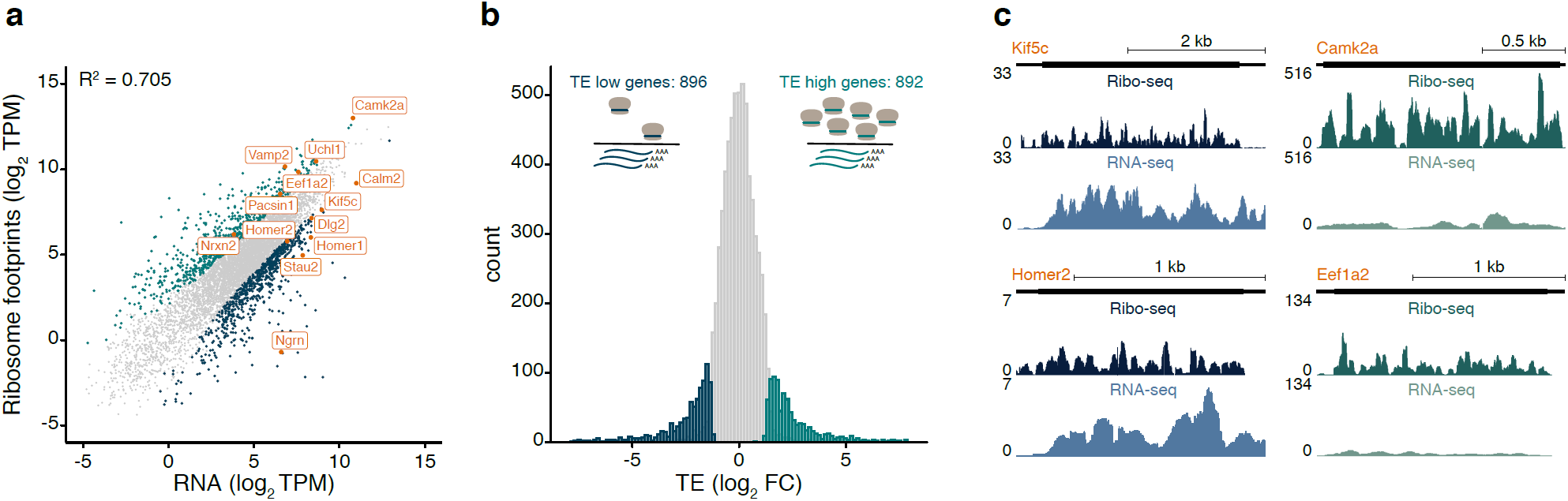
A substantial fraction of transcripts is regulated at the level of mRNA translation in synaptic regions. (**a**) Correlation of the local translatome with the local transcriptome. Scatter plot of the ribosome footprint versus mRNA levels (log_2_ TPM) in the neuropil (R^2^ = 0.705, p < 2.2e-16). Highlighted are genes with higher (teal) or lower (blue) ribosome footprints to mRNA ratios. (**b**) Histogram of the translational efficiency (TE) (log_2_ FC) in the neuropil, defined as the ratio of ribosome footprints from Ribo-seq to mRNA reads from RNA-seq (see Methods). Highlighted are genes with significantly higher (TE_high_, teal) or lower (TE_low_, blue) TE than 0.58 (FDR < 0.05). (**c**) Coverage tracks representing the average ribosome footprint (top) or RNA (bottom) coverage for candidate TE_low_ (*Kif5c, Homer2*) or TE_high_ (*Camk2a, Eef1a2*) genes in the neuropil. The y-axis indicates reads per million (RPM).

To understand how the transcript-specific translational control near synapses might be achieved, we extracted sequence features of neuropil TE_high_ and TE_low_ transcripts. We observed that, as a group, TE_low_ transcripts had longer coding sequences, consistent with previous observations^16,22-25^ (Fig. S4a). The efficiency of mRNA translation is also influenced by elements within the UTRs that serve as binding platforms for regulatory RBPs^11,12^. Because longer UTRs harbor more cis-acting elements^11,13^, we examined the 3’ and 5’UTR length of the neuropil translationally regulated transcripts. We found that TE_low_ genes exhibited significantly longer 5’ and 3’UTRs (Fig. 4a, b). To identify potential RBPs for the neuropil UTRs, we searched for known RBP consensus motifs^26^ and determined whether transcript groups sharing the same motifs were associated with higher or lower TE values (see Methods). Ninety-five 3’UTR motifs targeted by 43 RBPs were associated with transcripts displaying significantly higher TE values (Fig. 4c). For example, consistent with their described role as translational enhancers^27-29^ HNRNPK and MBNL1 motifs were detected in transcripts exhibiting significantly higher TE values (Fig. 4c). On the other hand, 119 3’UTR motifs targeted by 81 RBPs were associated with transcripts exhibiting significantly lower TE values (Fig. 4c). Among these, we identified, for example, the CPEB, Hu (Elav) and PUF/Pumilio RBP families, all known for their repressive action on translation in neuronal processes^30^. We note that none of the RBP motifs we detected within neuropil 5’UTRs were associated with transcripts displaying significantly higher or lower translation efficiencies (Fig. S4b). Our results thus reveal the identity of potentially novel regulators that bind the 3’UTR and control translation, either directly or indirectly (e.g. through the regulation of polyadenylation^26^ or mRNA decay^27^).

**Figure 4.**
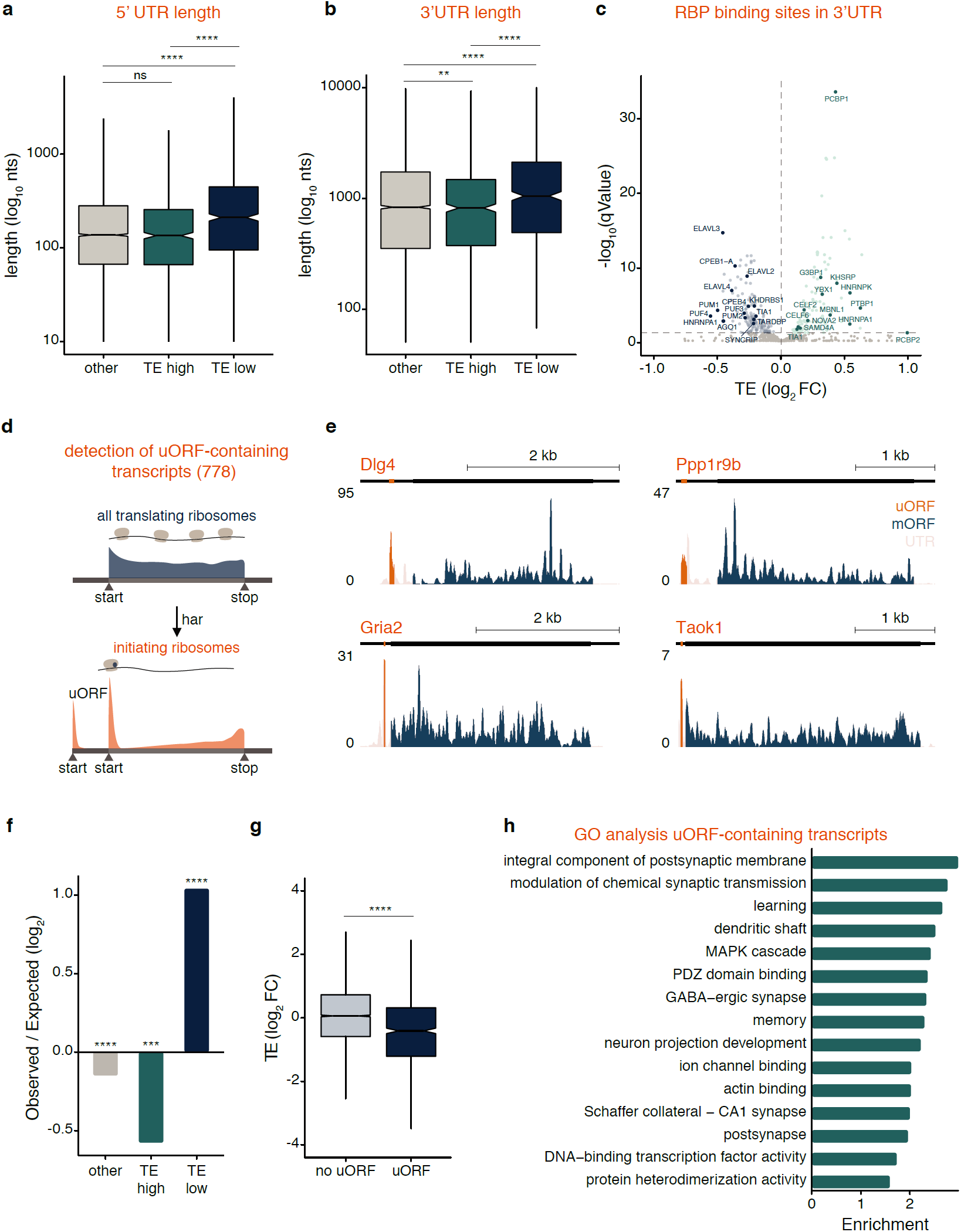
Features of translationally regulated transcripts in synaptic regions. (**a, b**) Box plots of 5’UTR (**a**) and 3’UTR (**b**) length (log_10_ nucleotides (nts) for TE_high_ (teal), TE_low_ (blue) and other (gray) genes. Bars indicate 1.5*IQR. **p < 0.01, **** p < 0.0001; one-way ANOVA test followed by pairwise t-test with BH p-value adjustment. (**c**) Shown are RNA-binding protein (RBP) motifs within 3’ UTRs associated with significantly lower (blue) or higher (teal) TE values (qValues < 0.05; Wilcoxon rank sum test; see Methods). (**d**) Detection of translated upstream open reading frames (uORFs) in hippocampal neurons. Translation initiation sites were mapped using the drug harringtonine (har), which accumulates ribosomes at start codons. A total of 778 uORF-containing neuronal transcripts were detected in the neuropil. (**e**) Coverage tracks representing the average ribosome footprint reads along the UTRs (beige), detected uORFs (orange) or the main protein coding sequence (blue) of *Dlg4, Gria2, Taok1* and *Ppp1r9b* in the neuropil. The y-axis indicates reads per million (RPM). (**f**) Observed-to-expected ratio of TE_high_ (teal), TE_low_ (blue) and other (gray) transcripts containing uORFs. ***p < 0.001, ****p < 0.0001; hypergeometric test. (**g**) Neuropil TE (log_2_ FC) measurements of transcripts containing translated uORFs (“uORF”) or not (“no uORF”). ****p < 0.0001; Welch two-sample t-test. (**h**) GO terms representing the top fifteen significantly (FDR < 0.05) enriched protein function groups for uORF-containing transcripts in the neuropil.

Upstream open reading frames (uORFs) play an important role in regulating the translation of the main protein coding sequence^31^. While most uORFs are believed to exert a negative effect on the translation of downstream ORFs^31^, a few examples of positive-acting uORFs have been reported^32,33^. To identify translated uORFs in the neuropil, we took advantage of the drug harringtonine, which causes the accumulation of ribosomes at start codons^34^ (Fig. 4d). After treating neurons with harringtonine (150 s), we mapped upstream translation initiation sites within neuronal transcripts (Fig. 4d). In total we identified 778 uORF-containing mRNAs in the neuropil (Fig. 4d, see Methods^35^) including novel (e.g. *Gria2, Toak1, Dlg4* and *Ppp1r9b* (Fig. 4e, Fig. S4c) and previously described (e.g. *Atf4* and *Ppp1r15b*^36,37^, Fig. S4d) transcripts. A comparison of TE_low_ and TE_high_ neuropil transcripts revealed an overrepresentation of uORF-containing transcripts in the TE_low_ group and an underrepresentation of uORF-containing transcripts in the TE_high_ group (Fig. 4f). Additionally, uORF-containing transcripts displayed a significantly lower neuropil median TE value when compared with non-uORF-containing mRNAs (Fig. 4g). Consistent with this, we found a weak negative correlation between the neuropil TE of uORF-containing transcripts and the uORF/mainORF (mORF) ribosome footprint ratio (Fig. S4e). Thus, for neuropil mRNAs, ribosome occupancy of uORFs is often associated with translational repression of the downstream “main” protein coding sequence of neuropil transcripts. A Gene ontology (GO) analysis indicated that above described uORF-containing neuropil mRNAs were significantly enriched for terms like “modulation of chemical synaptic transmission”, “learning” and “memory” (Fig. 4h). These findings highlight uORFs as an important translational regulatory element present in many synaptic transcripts.

### Many transcripts exhibit differential translation between neuronal compartments

An open question concerns the relative contribution of central (somata) and local (neuropil) sources to the total level of a given neuronal protein. To identify transcripts that exhibit differential translation between the somata and neuropil, we computed neuropil:somata Ribo-seq ratios (see Methods). We found ∼ 800 transcripts that exhibited significantly increased translation levels in the neuropil compared to the somata (“neuropil-translation-up”, Fig. 5a). These included, for example *Shank1, Map2* and *Dgkz* (Fig. 5a, b). In contrast, 2945 transcripts were more translated in the somata, including *Gria2, Neurod6* and *Hpca* (“somata-translation-up”, Fig. 5a, b).

**Figure 5.**
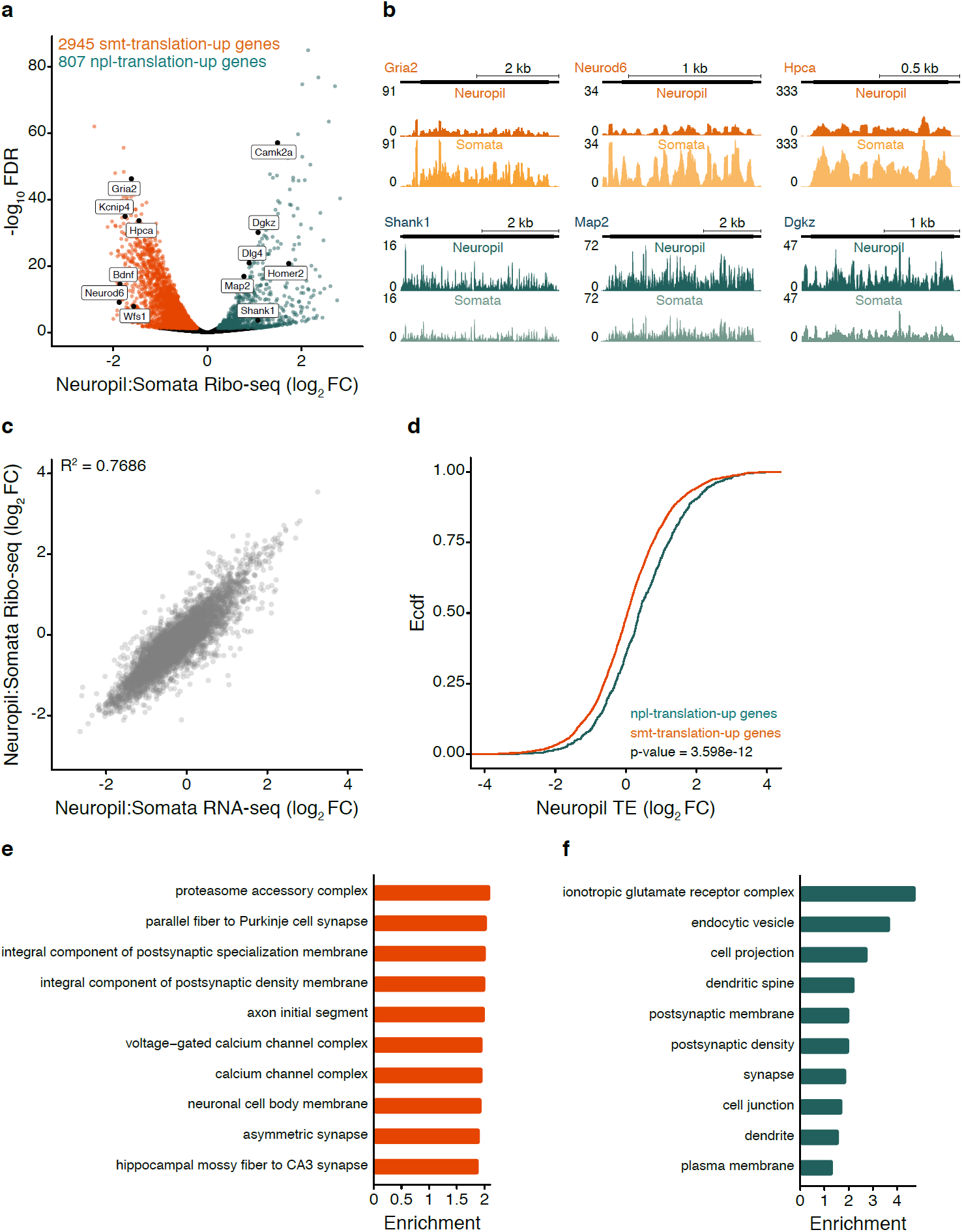
Many transcripts display differential translation between the somata and neuropil. (**a**) Volcano plot comparing the translational level of transcripts between compartments (neuropil:somata Ribo-seq ratio (log_2_ FC)). FDR < 0.05 using DESeq2 (see Methods). Colored dots highlight the transcripts significantly more translated in the somata (somata (smt)-translation-up, n = 2945, orange) or neuropil (neuropil (npl)-translation-up, n = 807, teal). (**b**) Coverage tracks representing the average neuropil (top) or somata (bottom) ribosome footprint coverage for candidate somata-translation-up (*Gria2, Neurod6, Hpca*) and neuropil-translation-up (*Shank1, Map2, Dgkz*) transcripts. The y-axis indicates the number of normalized reads. (**c**) Correlation of between-compartment transcriptional (RNA-seq neuropil:somata ratio (log_2_ FC), x-axis) and translational (Ribo-seq neuropil:somata ratio (log_2_ FC), y-axis) changes (R^2^ = 0.7686, p < 2.2e-16). (**d**) Empirical cumulative distribution frequency (Ecdf) of the TE (log_2_ FC) of somata-translation-up (orange) and neuropil-translation-up (teal) transcripts in the neuropil. p = 3.598e-12, Kolmogorov-Smirnov test. (**e, f**) GO terms representing the top 10 highest significantly enriched (FDR < 0.05) protein function groups for somata-translation-up (**e**) and neuropil-translation-up (**f**) transcripts.

We next asked how much the translational differences between the somata and neuropil might be explained by differences in the abundance of localized mRNAs by computing a neuropil:somata RNA-seq ratio. A comparison of the neuropil:somata Ribo-seq and RNA-seq ratios revealed that the translational changes between compartments correlated well (R^2^ = 0.766) with the differences in RNA abundance. Thus, many but not all of the between-compartment differences in ribosome footprint levels can potentially be accounted for by differences in the amount of mRNA present (Fig. 5c). Nevertheless, the neuropil-translation-up transcripts exhibited significantly higher TEs in both neuropil (Fig. 5d) and somata (Fig. S5). These findings indicate that the neuropil-translation-up mRNAs likely contain inherent RNA sequence features that favor their translation at higher efficiency in both compartments.

Previous studies suggested that neuronal mRNAs might be translationally silenced, via the “pausing” of ribosomes at the level of elongation, during their transport from the cell body^14,38^. To address this, we asked whether the neuropil- and somata-translation-up transcripts exhibited differences in the speed of translation elongation. We performed a time-series of ribosome “run-off” by incubating cultured hippocampal neurons for 15, 30, 45, or 90s with harringtonine, resulting in a progressive “run-off” of ribosomes over time (Fig. S6a, b). We analyzed the rate of ribosome progression from the 5’ ends of neuropil- and somata-translation-up transcripts (Fig. S6b). The neuropil- and somata-translation-up transcript subsets displayed a similar elongation rate of ∼ 4 codons per second (Fig. S6b, c), a value that agrees with measurements in other cell types (3-10 codons per second)^34,39-41^. Together, these findings indicate that enhanced efficiency of neuropil-translation-up transcripts likely results from regulation at the step of initiation but not elongation.

To examine whether particular protein function groups are encoded by transcripts that exhibit higher translation levels in either compartment, we performed a GO analysis. An enrichment of terms associated with synaptic function was found for both somata- and neuropil-translation-up transcripts (Fig. 5e, f). We observed a significant overrepresentation of the terms such as “proteasome accessory complex”, “axon initial segment” or “neuronal cell body membrane” for the somata-translation-up transcripts (Fig. 5e). On the other hand, mostly post-synaptic functions were significantly associated with the neuropil-translation-up transcripts, including for example “dendritic spine” and “post-synaptic density” (Fig. 5f). To understand better the synaptic function of the neuropil- and somata-translation-up transcripts, we analyzed the neuropil:somata Ribo-seq fold-changes of excitatory synaptic proteins (Fig. 6a). We noted that ionotropic and metabotropic glutamate receptor subunits (AMPARs, NMDARs and mGluRs) mostly displayed greater translation levels in cell-bodies (Fig. 6a). In contrast, many glutamate receptor-associated accessory (e.g. *Cnih2*) or scaffold proteins (e.g. *Shank1, Dlg4* and *Homer2*) were preferentially translated at synapses (Fig. 6a). Also, we found that the majority of pre-synaptic proteins exhibited greater protein synthesis rates in the somata (Fig. 6a). Lastly, we note that many of the neuropil-translation-up mRNAs are prominent on the gene hit-lists for neurodevelopmental and neurodegenerative disorders (Fig. 6b; see Fig. S7 for somata-translation-up transcripts) - suggesting that a focus on local regulatory mechanisms will be important for insights into both the pathology and the cure.

**Figure 6.**
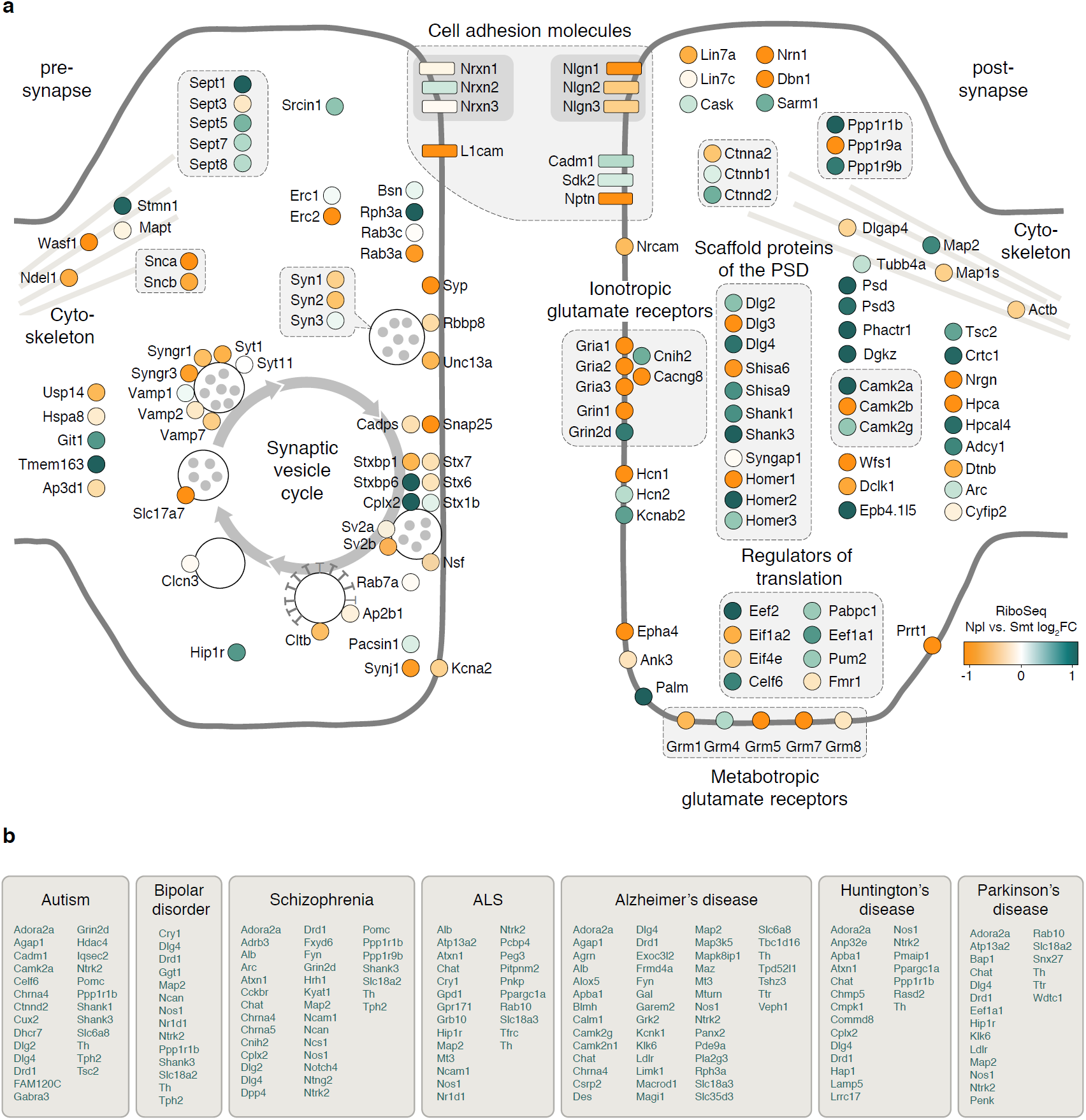
Functional segregation of transcripts differentially translated between the somata and neuropil. (**a**) Scheme depicting proteins of glutamatergic synapses. Ribo-seq neuropil:somata ratios (log_2_ FC) are color-coded from orange (more somata-translated) to teal (more neuropil-translated). Interacting proteins are displayed in closer proximity. Proteins with similar functions are grouped together and the synaptic vesicle cycle is indicated by arrows. (**b**) Shown are neuropil-translation-up genes associated with each of the neurodevelopmental and neurode-generative diseases. Some genes are listed for multiple diseases.

Using ribosome profiling, we detected more than 4800 mRNA species that are translated in synaptic regions, dramatically expanding the contribution of the local protein synthesis to the protein pool detected in dendrites, axons or synapses^42-45^. Indeed, among the locally translated mRNAs, we identified most protein families including signaling molecules (kinases or phosphatases), ion channels, metabotropic and ionotropic receptors, cell adhesion molecules, scaffold proteins as well as regulators of cytoskeleton remodeling or translation.

We detected a wide-spread translational regulation in synaptic regions under basal conditions, with an unexpectedly high dynamic range in the translation efficiencies of localized transcripts. Among the mechanisms that regulate local synthesis of synaptic proteins, we identified uORF-mediated translational control. This mode of translational control is often fine-tuned by the phosphorylation of eukaryotic initiation factor 2α (eIF2α)^46^. The phosphorylation of eIF2α inhibits global translation whilst leading to a paradoxical increase in the translation of a subset of uORF-bearing transcripts^47^. Many manipulations of synaptic activity modulate the phosphorylation status of eIF2α in neurons in vivo and in vitro^47-49^. Thus, activity-driven eIF2α phosphorylation could act as a switch to enhance the local translational efficiency of uORF-containing transcripts encoding key plasticity-related proteins.

Many transcripts were found differentially translated between neuronal compartments, with more than 800 mRNAs displaying enhanced translation levels in the neuropil. For many, the abundance of the mRNA was positively associated with the translation level differences between somata and neuropil, as observed previously in developing neurons derived from mouse embryonic stem cells^50^. Nevertheless, we also found that neuropil-translation-up transcripts were translated at higher efficiencies compared to somata-translation-up mRNAs. The positive translational regulation of neuropil-translation-up mRNAs was independent of their subcellular localization, indicating that this feature is intrinsic to the transcripts rather than influenced by the environment. Notably, the neuropil-translation-up transcripts often encoded signaling and scaffold proteins that play an important role in the maintenance and modification of synaptic strength. Thus, the translational enhancement of this transcript subset likely provides a means to ensure the efficient production of key synaptic proteins at very remote locations from the cell body where fewer mRNA copy numbers^4^ and ribosomes^16^ have been detected. Taken together, our results reveal an enormous capacity for local protein synthesis at hippocampal synapses in vivo. The findings overturn the prevailing belief that local translation is only a minor source of synaptic proteins and is only used to remodel the local proteome during synaptic plasticity^2^.

## Methods

### Animals

Timed pregnant SPF (Charles River Laboratories) females were housed in the institute’s animal facility for 1 week on a 12/12-hour light/dark cycle with food and water ad libitum until the litter was born. Cultured neurons were derived from P0 (postnatal day 0) Sprague-Dawley rat pups (CD Crl:CD, both male and female, RRID: RGD 734476). Pups were euthanized by decapitation. The housing and euthanasia procedures involving animal treatment and care were conducted in conformity with the institutional guidelines that are in compliance with national and international laws and policies (DIRECTIVE 2010/63/EU; German animal welfare law; FELASA guidelines). The animals were euthanized according to annex 2 of § 2 Abs. 2 Tierschutz-Versuchstier-Verordnung. Animal numbers were reported to the local authority (Regierungspräsidium Darmstadt, approval numbers: V54-19c20/15-F126/1020 and V54-19c20/15-F126/1023).

### Cell culture

#### Primary hippocampal cultures

Dissociated rat hippocampal neurons were prepared from P0 day-old rat pups as described previously^51^. <u>For the harringtonine treatment:</u> Hippocampal neurons were plated at a density of 31,250 cells/cm^2^ onto poly-d-lysine-coated 100 mm dishes and cultured in pre-conditioned growth medium (Neurobasal-A, B27, GlutaMAX, 30 % glia-culture supernatant, 15 % cortex-culture supernatant) for 21 days in vitro (DIV). At one DIV, cells were treated with Ara-C (Sigma) at a final concentration of 5 µM to prevent the overgrowth of non-neuronal cells. After 48 h, medium was replaced with pre-conditioned growth medium and cells were cultured until 21 DIV. Cells were fed with one ml of pre-conditioned medium every 7 days. Three independent biological replicates were prepared. <u>For the in situ visualization of newly synthesized proteins</u>: 30,000 hippocampal cells were plated onto poly-d-lysine coated glass-bottom Petri dishes (MatTek) and cultured until 18 DIV. All primary cultures were maintained in a humidified incubator at 37 °C and 5 % CO_2_.

#### Neuron-enriched and glia-enriched cultures

Neuron-enriched and glia-enriched cultures were prepared from the same litter as described previously^8^. The hippocampi of P0 day-old rat pups were isolated and triturated after digestion with papain. Both cultures were plated on 60 mm cell culture dishes. For the preparation of hippocampal neuron-enriched cultures, cells were plated onto poly-d-lysine-coated 60 mm cell culture dishes and treated as described above with Ara-C (Sigma) at a final concentration of 5 µM for 48 h. After 48 h, medium was replaced with pre-conditioned growth medium and cells were cultured until 21 DIV. For the preparation of glia-enriched cultures, cells were plated onto uncoated 60 mm cell culture dishes in conditioned minimal essential medium (Minimal essential medium, 10 % horse serum, 0.6 % glucose (w/v)). At 7 DIV, medium was replaced with pre-conditioned growth medium and cells were cultured until 21 DIV. Four independent biological replicates were prepared.

### Harringtonine treatment of hippocampal cultures

At 24 h before drug treatment, cell medium was adjusted to 8 ml per dish. Harringtonine (LKT Laboratories) was added to a final concentration of 2 μg/ml from a 5 mg/ml stock in 100% ethanol. Cells were returned to the incubator at 37 °C for 15, 30, 45, 90 or 150 s. Cycloheximide was added to a final concentration of 100 μg/ml from a stock of 50 mg/ml in 100% ethanol. After drug addition, cells were returned to the incubator at 37°C for 1 min. After the incubation with cycloheximide, the cells were immediately placed on ice and washed twice with ice-cold PBS plus 100 μg/ml cycloheximide and lysed in polysome lysis buffer as described below. Ribo-seq libraries were prepared as described below. The 0, 30 and 90 s datasets were previously published in^16^.

### Direct visualization of newly synthesized proteins in situ

Nascent polypeptides were visualized as described previously^20^. In brief, primary hippocampal cultures were incubated for 5 min with puromycin at a final concentration of 3 µM or with an equal volume of water as vehicle control. For the protein synthesis inhibitor control, cells were pre-incubated for 30 min with anisomycin at a final concentration of 40 µM. Cells were washed twice with pre-warmed growth medium and fixed for 20 min in PFA-sucrose (4 % paraformaldehyde, 4 % sucrose, 1 mM MgCl_2_, 0.1 mM CaCl_2_ in 1X PBS). Fixed cells were washed twice in 1X PBS, permeabilized for 15 min in 0.5 % Triton X-100 in blocking buffer (4 % goat serum in 1X PBS) and blocked for one hour in blocking buffer at room temperature. Primary antibodies against the protein-of-interest, against puromycin, and against MAP2 were diluted in blocking buffer and incubated for 90 min at room temperature. Cells were washed three times in 1X PBS and then incubated with the appropriate PLA-antibodies (PLA^minus^, PLA^plus^, Duolink, 1:10) diluted in blocking buffer and a secondary antibody to visualize MAP2 (A488, ThermoFisher, A-11073, 1:1000) for one hour in a pre-warmed humidified chamber at 37 °C. Cells were washed 5 times with wash buffer A (0.01 M Tris, 0.15 M NaCl, 0.05 % Tween 20) and incubated for 30 min with the ligation reaction including the circularization oligos and the T4 ligase prepared according to the manufacturer’s recommendations in a pre-warmed humidified chamber at 37 °C. Cells were washed three times in wash buffer A and incubated in the amplification reaction mixture containing Phi29 polymerase and the fluorophore-labeled detection oligo prepared following the manufacturer’s recommendations in a pre-warmed humidified chamber for 100 min at 37 °C. The amplification was stopped by three washes in wash buffer B (0.2 M Tris, 0.1 M NaCl, pH 7.5) and 5 washes in 1X PBS. The cells were imaged immediately in 1X PBS. Fluorescence imaging was performed with an LSM780 confocal microscope (Zeiss) using a 40x objective (Plan-Apochromat 40x/1.4 Oil DIC M27) and 0.6x digital zoom. Appropriate excitation laser lines and spectral detection windows were used. Laser power and detector gain were adjusted to avoid saturated pixels. Imaging conditions were held constant between conditions. Z-stacks, spanning the entire cell-width, were obtained. For visualization, z-stacks were projected as maximum intensity projections and dendrites were blindly selected to be straightened. The brightness and contrast of the Puro-PLA signal was first adjusted, then binarized and finally dilated for better visualization. Two independent experiments with at least two dishes per condition were performed.

### Preparation of the RNA-seq libraries

For the neuron- and glia-enriched cultures, RNA was isolated from lysates using the Direct-zol RNA micro prep kit (Zymo). RNA integrity was assessed using the Agilent RNA 6000 Nano kit. mRNA-seq libraries were prepared starting from ∼200 ng of total RNA, using the TruSeq stranded mRNA library prep kit (Illumina). Libraries were sequenced on an Illumina NextSeq500, using a single-end, 75-bp run. Rat CA1 somata and neuropil total RNA-seq libraries were previously published in^16^.

### Preparation of the Ribo-seq libraries

Ribo-seq libraries from hippocampal neuron cultures treated for 15, 45 and 150 s with harringtonine were prepared as described previously^16^. The 0, 30 and 90 s datasets were previously published in^16^.

### Translating ribosome affinity purification (TRAP)

The input- and TRAP-seq datasets from hippocampi of Camk2a-Cre-RiboTag or somata / neuropil sections of Wfs1-Cre-RiboTag mice were previously published in^16^.

### Data Analysis

#### Genome and transcriptome alignment of ribosome profiling libraries

Sequencing adapters were trimmed using the Cutadapt software version 1.15 ^52^ with the following arguments: *--cut 1 --minimum-length 22 --discard-untrimmed --overlap 3 -e 0*.*2*. An extended unique molecular identifier (UMI) was constructed from the two random nucleotides of the reverse transcription primer and the five random nucleotides of the linker and added to the FASTQ description line using a custom Perl script. To remove reads originating from non-coding RNA (ncRNA, i.e. rRNA), trimmed reads were aligned to rat ncRNA using Bowtie2 version 2.3.5.1 (--very-sensitive)^53^ and aligned reads were discarded. The remaining reads were aligned to the rat genome (rn6) with the split-aware aligner STAR version 2.7.3.a^54^ with the following arguments: *--twopassMode Basic --twopass1readsN -1 -- seedSearchStartLmax 15 --outSJfilterOverhangMin 15 8 8 8 -- outFilterMismatchNoverReadLmax 0*.*1*. To retrieve transcript coordinates, STAR’s quant mode (--quantMode) was used. Throughout the study, genome alignments were used for differential expression analyses and genomic feature analyses. Transcriptome alignments were used for all other analyses. The STAR genome index was built using annotation downloaded from the UCSC table browser^55^. PCR duplicates were suppressed using a custom Perl script and alignments flagged as secondary alignment were discarded before analysis. Only footprints with sizes between 24 and 34 nts were used for analyses.

### Genome alignment of RNA libraries

Sequencing adapters and low-quality nucleotides were trimmed using the Cutadapt software version 1.15^52^ with the following arguments: *--minimum-length 25 --nextseq-trim=20*. The trimmed reads were aligned to the rat (rn6) or the mouse (mm10) genome with STAR version 2.7.3a^54^.

### Genomic feature analysis

The coordinates of genomic features (CDS, 3’UTR, 5’UTR, intron) were downloaded from the UCSC table browser in BED format^55^. Bedtools version 2.26.0^56^ was used to convert BAM into BED files and to identify reads overlapping with the individual features.

### Three-nucleotide periodicity

P-site offsets were defined for different footprint lengths. Each footprint start position defined the footprint frame in reference to the annotated start codon. The footprint reads were virtually back projected over the start codon and the offsets from the start and the end of the read were calculated. We used every read of a given length and accumulated the most probable offset and frame. Next, the P-site position per footprint read was deduced from its length and the previously determined offset. All P-site positions were plotted for 100 nucleotides around the start-, the stop-codon and the center of a transcript. To correct for differences in translation rates between genes, the P-site coverage of each gene was normalized to its mean footprint coverage. The nucleotide coverage at the 0, 1 and 2 frame positions were assessed. A one-way analysis of variance (ANOVA) was used to determine if the observed frame fraction was different from the expected frame fraction. A significant p-value rejected the null-hypothesis that all frames featured the expected P-site coverage.

### Genome browser track visualization

Footprint alignments were converted into the BedGraph file format using Bedtools version 2.26.0 and visualized as custom tracks on the UCSC Genome Browser^57^. Footprint coverages were corrected for sequencing depth.

### Differential expression analysis

#### RNA-seq and Ribo-seq neuropil:somata ratios

For both, total RNA sequencing and ribosome footprint libraries from the somata and neuropil, the software featureCounts version 2.0.0^58^ was used to calculate counts per gene from reads that were aligned to the rat genome. All annotated transcript isoforms were considered. Raw counts were fed into DESeq2 version 1.26.0 and LFC shrinkage was used^59^.

#### RiboTag IP:input ratios and neuron-enriched:glia-enriched culture ratios

The software featureCounts version 2.0.0^58^ was used to calculate counts per gene from reads mapped to the genome (mm10, rn6). All annotated transcript isoforms were considered. Raw counts were fed into DESeq2 version 1.26.0 and LFC shrinkage was used^59^.

### Gene ontology analysis

Gene ontology analysis was performed for neuropil- and somata-translation-up genes. All detected genes, without the contaminants, were used as background. Only GO terms with at least 10 hits in the background set were considered. Enrichment scores were calculated and the top10 GO terms of the category “cellular components” with an FDR below 0.05 were visualized.

Gene ontology analysis was performed for uORF-containing transcripts. All detected genes in the neuropil, without the contaminants, were used as background. Only GO terms with at least 50 hits in the background set were considered. Enrichment scores were calculated and the top5 GO terms for the categories “cellular component”, “biological process”, and “molecular function” with an FDR below 0.05 were visualized. For both analyses the in-house developed tool “GOAnalysis” was used^13^.

### Neurological disease-association

The neuropil- and somata-translation-up genes were analyzed for their association with neurodevelopemental and neurodegenerative disorders using GeneAnalytics (“MalaCards”, Lifemap Sciences)^60^.

### Motif analysis 3’and 5’UTR

RNA-binding protein (RBP) motifs (human, rat, and mouse) were downloaded as position weighted matrices from the public ATtRACT database^26^. The FIMO tool from the MEME suite version 5.1.1 was used to scan 5’ and 3’UTRs for motif occurrences, using the default threshold (p-value = 1e-4) and a pre-calculated nucleotide background model derived from query sequences^61^. Only genes with an RBP motif occurrence were considered for analysis. For each identified RBP motif, the motif-containing genes were grouped and a median translational efficiency (TE) value was calculated. A Wilcoxon rank sum test was conducted to test if the median TE of a given RBP motif group differed from the median TE of all genes that do not contain the motif.

### Detection of translated uORFs

The ORF-RATER pipeline (https://github.com/alexfields/ORF-RATER) was run as previously described^35^, starting with the harringtonine 150 s as well as the neuropil and somata BAM files. Note, that it is possible that a translated uORF may be assigned a low score, as ORF-RATER is tuned to indicate the highest-confidence sites of translation, at the expense of an increased false negative rate^62^. The following parameters were used: “--codons NTG” for ORF types, “--minrdlen 28 --maxrdlen 34” for harringtonine-treated samples, “-- minrdlen 27 --maxrdlen 34” for neuropil and somata samples. Only uORFs with a score of at least 0.7, a length of at least 3 codons and at least one count in each of the neuropil replicates were considered.

### Transcript feature analysis

5’ and 3’UTR lengths were calculated based on the Rattus Norvegicus annotation version 6 (rn6). 3’UTR lengths were corrected in accordance to newly identified 3’UTR isoforms described in^13^. For genes with multiple 5’UTR isoforms the longest 5’UTR sequence was chosen, giving priority to curated isoforms. For genes with multiple 3’UTRs the most expressed 3’UTR isoform was chosen^13^.

For the comparison of 5’UTR lengths between “TE_high_”, “TE_low_” and “others” only 5’UTRs with a minimum length of 10 nts and a maximum length of 5,000 nts were considered. For the comparison of 3’UTR lengths between “TE_high_”, “TE_low_” and “others” only 3’UTRs with a minimum length of 50 nts and a maximum length of 10,000 nts were considered.

### Metagene analysis and computation of the elongation rate

The coverage of each gene was projected along the CDS in transcript coordinates (only exons). Genes with CDS lengths shorter than 440 codons were omitted from analysis. Each metagene profile was scaled by the average coverage between codon 400 to 20 codons before the stop codon. For each time point, the metagene profiles were smoothed with a running average window of 30 codons. For each group, the coverage tracks were accumulated, averaged and normalized to the 0 s condition. A baseline coverage track was defined as 85% of the non-treated sample coverage track. The first positive crossing between the harringtonine-treated coverage track and the baseline coverage track determined the crossing position in codons. Elongation rates were calculated as the slope of a linear regression between the harringtonine incubation times for each track and the crossing position in codons.

### Computation of translational efficiency

The number of ribosomes per transcript was estimated by integrating Ribo-seq and RNA-seq libraries to calculate translational efficiency (TE) values in the neuropil. Raw Ribo-seq and RNA-seq counts, falling into gene coding sequences, were fed into DESeq2 version 1.26.0 and LFC shrinkage was used^59^. TE values that were either significantly higher than log_2_(1.5) or smaller than log_2_(1.5) were assigned to TE_high_ and TE_low_, respectively (lfcThreshold = log_2_(1.5) with an adjusted p-value < 0.05).

### Statistical analyses

Additional information about statistical analyses not specified in the figure legends.

Figure 4a: One-way ANOVA: F-value = 33.43; p-value = 3.55e-15

Figure 4b: One-way ANOVA: F-value = 15.73; p-value = 1.54e-07

Figure 4g: Welch Two Sample t-test: t = -10.093; df = 983.29; p-value < 2.2e-16; uorf-mean = -0.4096, no uorf-mean = 0.0691; two-sided

Figure 5d: Two-sample Kolmogorov-Smirnov test: D = 0.151; p-value = 3.598e-12; two-sided

Figure S1e: One-way ANOVA: F-value = 17.69; p-value = 0.0031

Figure S1f: One-way ANOVA: F-value = 13.49; p-value = 0.006

Figure S4a: One-way ANOVA: F-value: 98.48; p-value < 2e-16

Figure S5: Two-sample Kolmogorov-Smirnov test: D = 0.10388; p-value = 5.526e-06; two-sided

Figure S6c: ANOVA: F-value = 0.5671; p-value = 0.5738

## Data and software availability

The accession number for the raw sequencing data published previously in^16^ is NCBI BioProject: PRJNA550323. All bioinformatic tools used in this study are contained in one modular C++ program called RiboTools. The source code and further notes on the algorithms can be found on our GitHub repository doi:10.5281/zenodo.3579508. Other analysis scripts and codes are available upon request.

## Acknowledgements

We thank Elena Ciirdaeva for help with mRNA library preparation. A.B. is supported by an EMBO long-term postdoctoral fellowship (EMBO ALTF 331-2017). E.M.S. is funded by the Max Planck Society, an Advanced Investigator award from the European Research Council (grant 743216), DFG CRC 1080: Molecular and Cellular Mechanisms of Neural Homeostasis, and DFG CRC 902: Molecular Principles of RNA-based Regulation.

## Contributions

C.G. and A.B. designed and conducted experiments and analyzed results. G.T. analyzed results. I.B., B.N. and S.T.D. conducted experiments. E.M.S. designed experiments and supervised the project. C.G., A.B. and E.M.S. wrote the manuscript, and all authors edited the manuscript.

## Correspondence

Correspondence to Erin M Schuman

---

**Figure S1.**
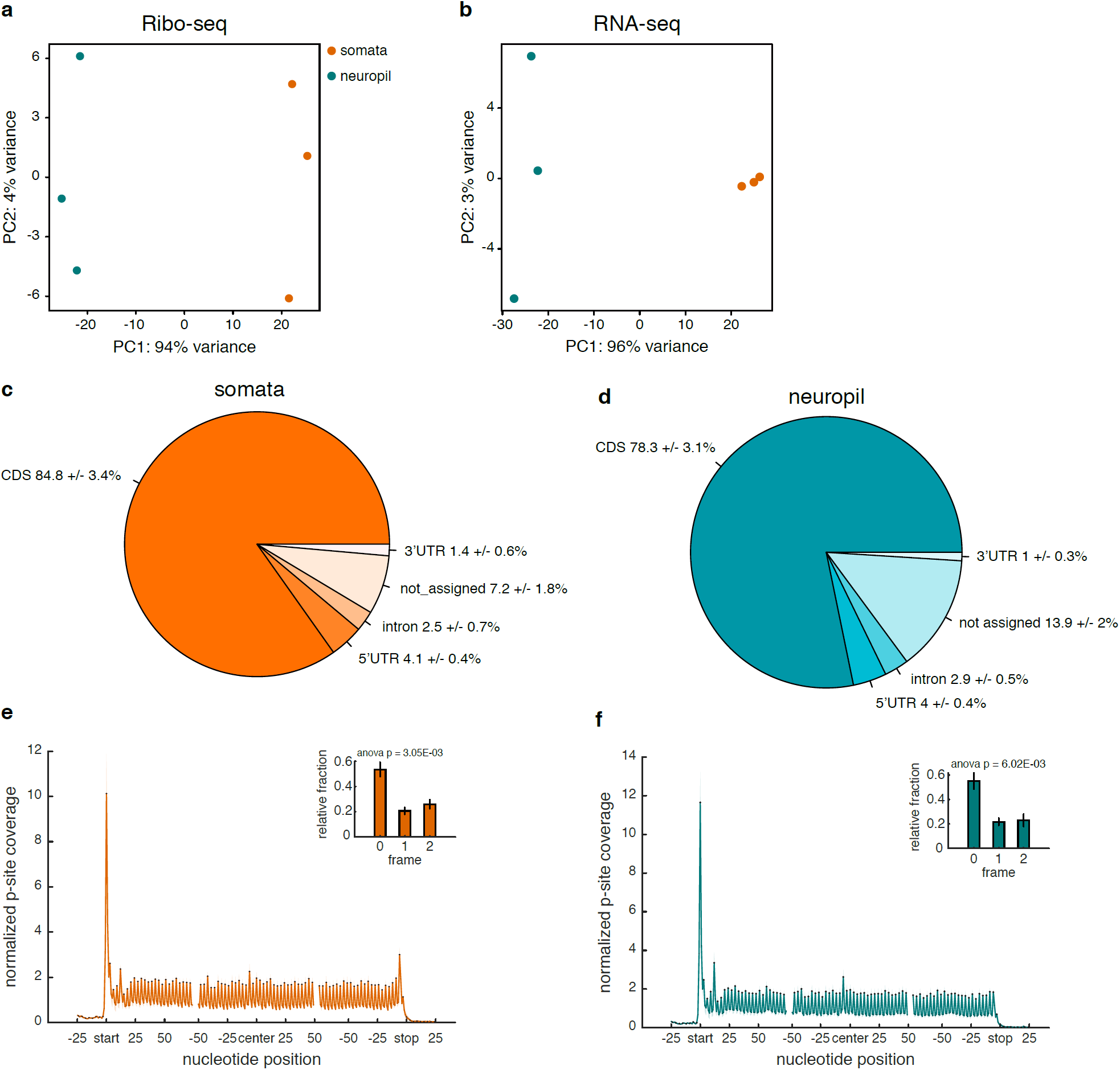
Quality metrics of the RNA- and Ribo-seq data. (**a, b**) Principal component analysis (PCA) showing the within and between group variance of neuropil and somata Ribo-seq (**a**) and RNA-seq (**b**) samples. (**c, d**) Percentage of ribosomal footprints of the somata (**c**) and neuropil (**d**) libraries, mapping to different genomic regions, including the coding sequence (CDS), 5’-/3’ untranslated regions (UTR) and introns. (**e, f**) Metagene analyses showing the P-site coverage of ribosomal footprints in the somata (**e**) and neuropil (**f**) samples. The mean normalized coverage is projected per nucleotide position around the start codon (start), central portion (center) and stop codon (stop) of the CDS. The standard error is shaded (n = 3). Bars in the insets indicate standard errors.

**Figure S2.**
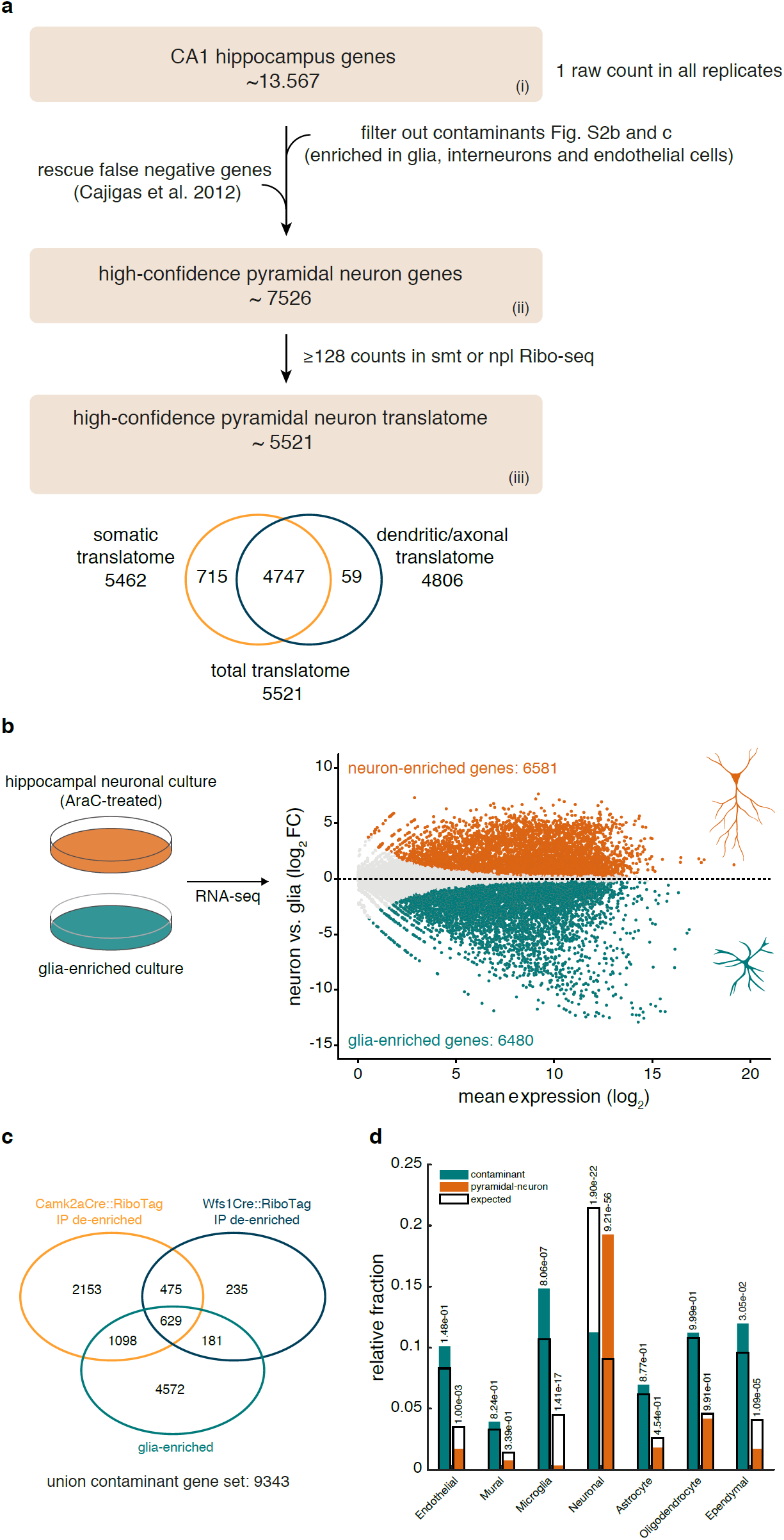
Bioinformatic filtering for neuronal transcripts. (**a**) Overview of the bioinformatic filtering steps, beginning with the removal of genes originating from non-excitatory neurons. Genes with an experimentally validated dendritic localization were rescued from the filter^4^. To define a high-confidence set of transcripts translated in excitatory neurons, a threshold of 128 mean counts across somata or neuropil replicates was chosen as a point where counting statistics contributed little to the inter-replicate variation^17^. (**b**) Identification of glia-enriched genes. Scheme outlining RNA sequencing in hippocampal neuron- and glia-enriched cultures. Differential expression analysis revealed 6581 and 6480 genes significantly enriched in neurons (orange) or glial (teal) cells, respectively. FDR < 0.01. (**c**) Venn diagram comparing the genes significantly de-enriched in the Rpl22-HA IP from the hippocampi of Camk2a-Cre-, the microdissected somata and neuropil layers of Wfs1-Cre-RiboTag mice^16^ and glia-enriched genes. The union of these datasets represents the list of “contaminant” non-excitatory neuron genes. (**d**) Observed-to-expected ratio of contaminant (teal) or excitatory neuron (orange) genes defined as described in (**a**) and previously associated with different hippocampal cell types by a single-cell study^18^. Numbers above the bars indicate the p-values for the enrichment (using a hypergeometric test).

**Figure S3.**
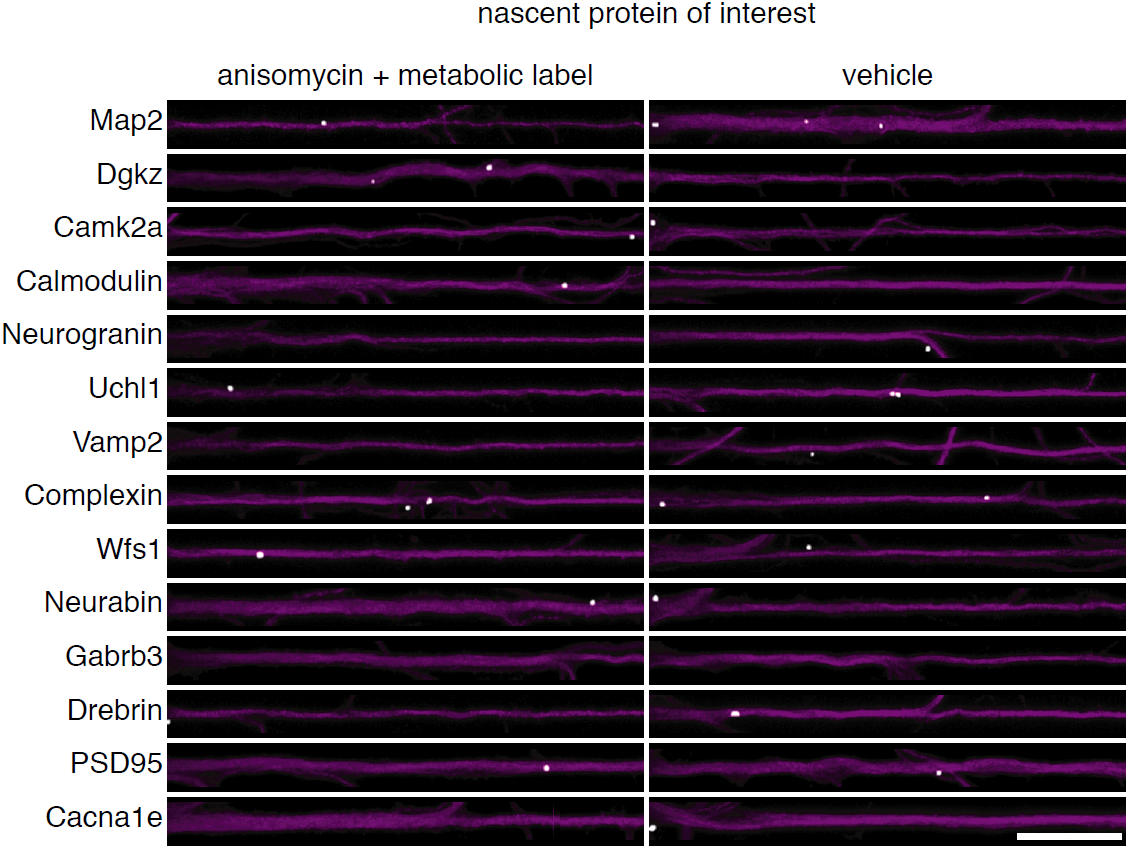
Specificity of the high-resolution fluorescent in situ detection of translation events in synaptic regions from cultured hippocampal neurons. Puro-PLA signal of candidate proteins of interest (white) in individual straightened dendrites from cultured hippocampal neurons in which the metabolic label was omitted (right) or neurons were pre-treated with the protein synthesis inhibitor anisomycin (left). Signal was dilated for better visualization. Anti-MAP2 immunolabeling of dendrites is indicated in magenta. Scale bar = 20 µm.

**Figure S4.**
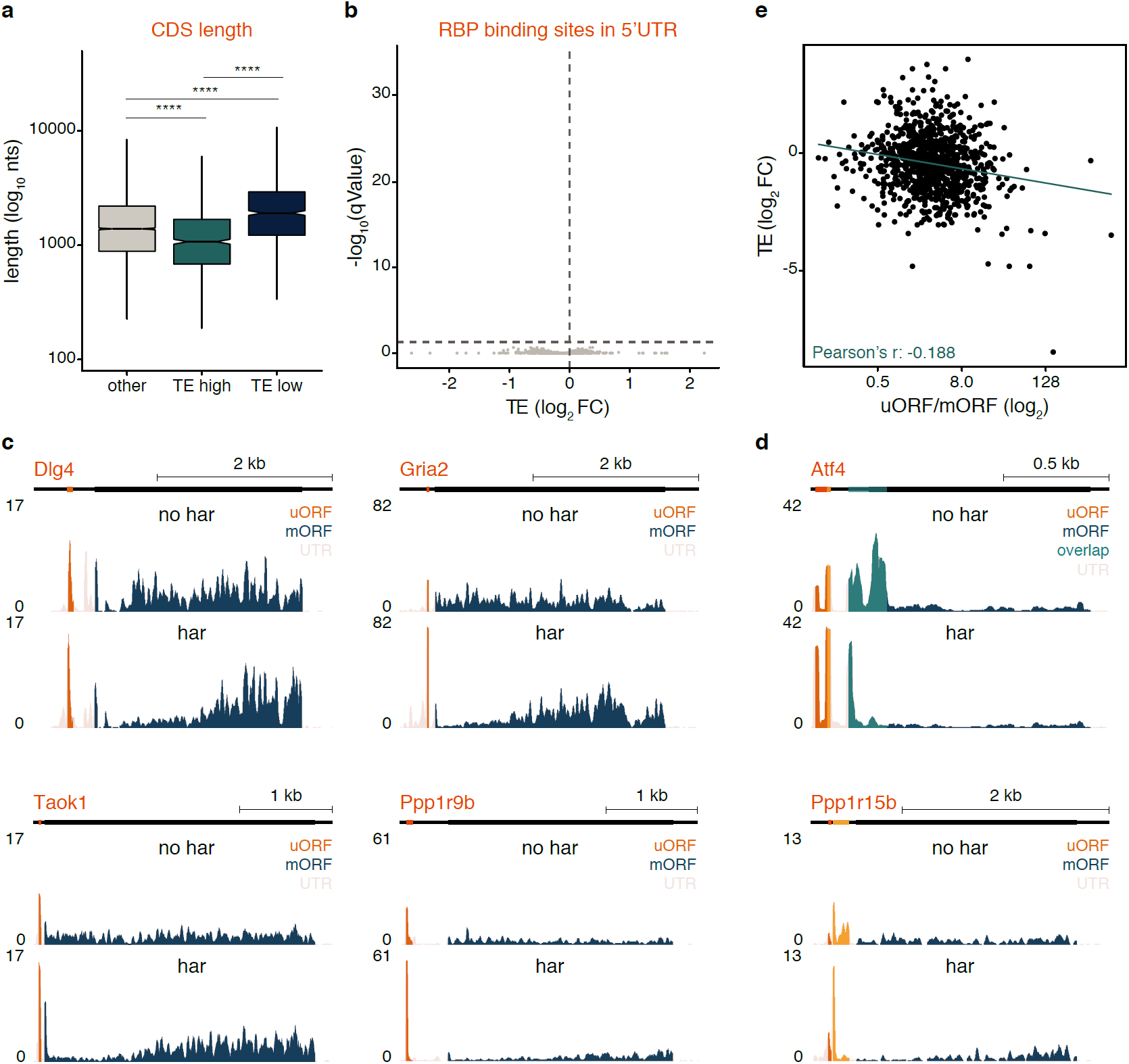
Features of translationally regulated transcripts in the neuropil. (**a**) Box plots of coding sequence (CDS) length (log_10_ nucleotides (nts)) for TE_high_ (teal), TE_low_ (blue) and other (gray) genes. Bars indicate 1.5*IQR. **** p < 0.0001; one-way ANOVA test followed by pairwise t-test with BH p-value adjustment. (**b**) No RNA-binding protein (RBP) motifs within 5’ UTRs were found significantly associated with lower or higher TE values (qValue < 0.05; Wilcoxon rank sum test; see Methods). (**c, d**) Coverage tracks representing the average ribosome footprint reads along UTRs (beige), uORFs (orange), overlapping uORFs (teal) or the main protein coding sequence (blue) in untreated (no har) or harringtonine-treated (har) primary hippocampal neuron cultures. The harringtonine-induced ribosome footprint accumulation at upstream start sites enabled the detection of novel uORFs within *Dlg4, Gria2, Taok1* and *Ppp1r9b* (**c**) and confirmed the well-studied uORF usage within the transcripts *Atf4* and *Ppp1r15b* (**d**). The y-axis indicates reads per million (RPM). (**e**) Neuropil TE measurements (log_2_ FC) are plotted as a function of uORF:mORF ribosome footprint ratio.

**Figure S5.**
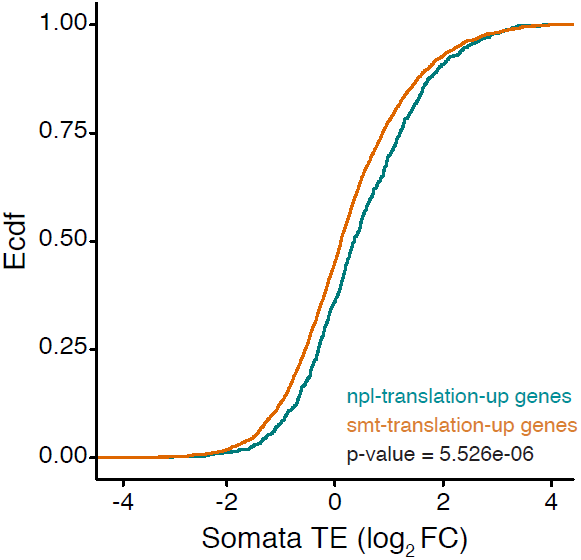
Neuropil-translation-up transcripts are translated at higher efficiency in the somata. Empirical cumulative distribution frequency (Ecdf) of the TE (log_2_ FC) of somata (smt)-translation-up and neuropil (npl)-translation-up genes in the somata. p = 5.526e-06, Kolmogorov-Smirnov test.

**Figure S6.**
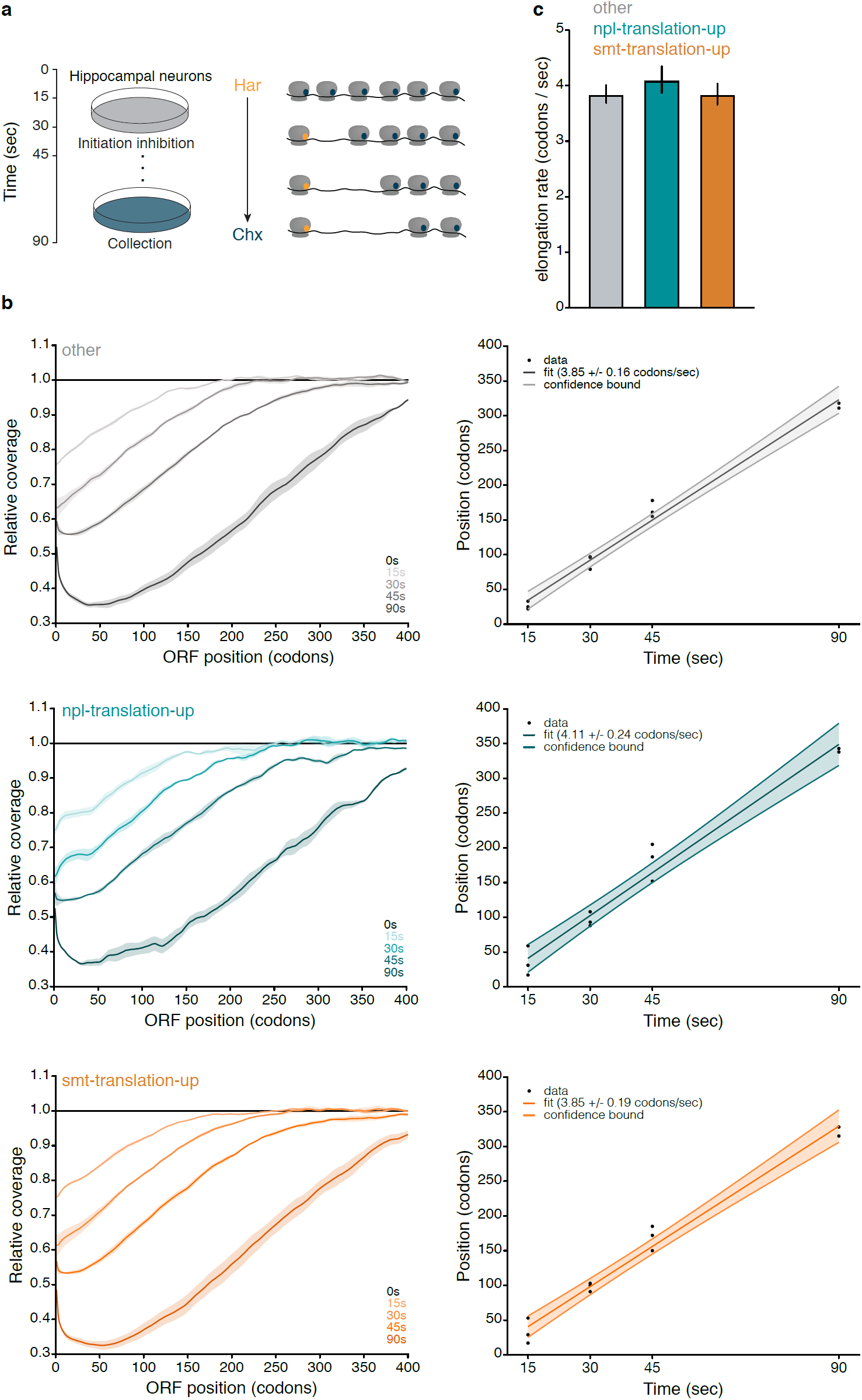
Neuropil- and somata-translation-up transcripts exhibit similar translation elongation rates. (**a**) Schematic depicting in vivo ribosome run-off following harringtonine incubation of rat hippocampal cultures. (**b**) Progressive loss of ribosome footprints from the 5’ end of somata (smt)-translation-up (orange), neuropil (npl)-translation-up (teal) and other (gray) transcripts. Ribosome densities of the samples treated with harringtonine for 15, 30, 45 and 90 s were normalized to the untreated control (n = 3). The codon position of 15% ribosome depletion was plotted as a function of harringtonine incubation times for smt-translation-up (orange), npl-translation-up (teal) and other (gray) transcripts. A linear fit was computed for each transcript subset. The standard errors are shaded. (**c**) Elongation rates for smt-translation-up (orange), npl-translation-up (teal) and other (gray) transcripts inferred from the slope of the linear fit shown in (**b**) are plotted with their standard error (n = 3). p = 0.5738, ANOVA. Har, harringtonine; Chx, cycloheximide.

**Figure S7.**
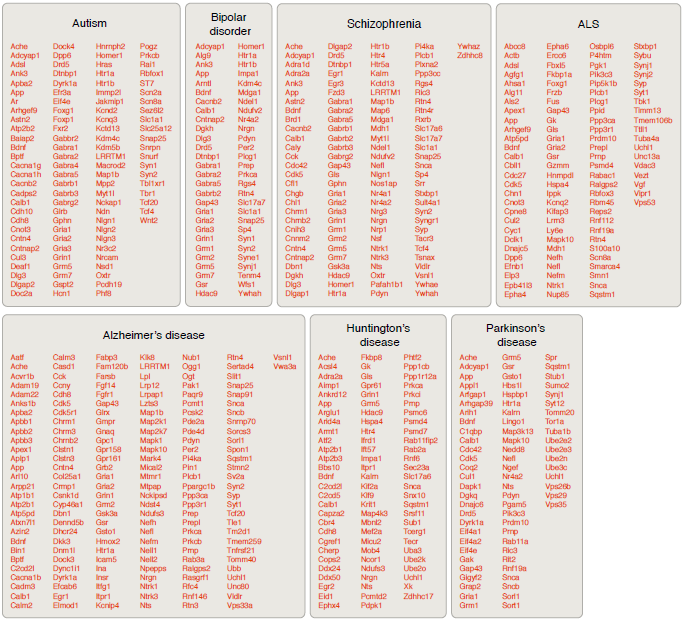
Neurological disease-associated transcripts with enhanced somatic translation. Shown are somata-translation-up genes associated with each of the neurodevelopmental and neurodegenerative diseases. Some genes are listed for multiple diseases.

